# TAF8 regions important for TFIID lobe B assembly, or for TAF2 interactions, are required for embryonic stem cell survival

**DOI:** 10.1101/2021.07.06.451281

**Authors:** Elisabeth Scheer, Jie Luo, Frank Ruffenach, Jean-Marie Garnier, Isabelle Kolb-Cheynel, Kapil Gupta, Imre Berger, Jeff Ranish, László Tora

## Abstract

The human general transcription factor TFIID is composed of the TATA-binding protein (TBP) and 13 TBP-associated factors (TAFs). In eukaryotic cells, TFIID is thought to nucleate RNA polymerase II (Pol II) preinitiation complex formation on all protein coding gene promoters and thus, be crucial for Pol II transcription. TFIID is composed of three lobes, named A, B and C. Structural studies showed that TAF8 forms a histone fold pair with TAF10 in lobe B and participates in connecting lobe B to lobe C. In the present study, we have investigated the requirement of the different regions of TAF8 for *in vitro* TFIID assembly, and the importance of certain TAF8 regions for mouse embryonic stem cell (ESC) viability. We have identified a TAF8 region, different from the histone fold domain of TAF8, important for assembling with the 5TAF core complex in lobe B, and four regions of TAF8 each individually required for interacting with TAF2 in lobe C. Moreover, we show that the 5TAF core-interacting TAF8 domain, and the proline rich domain of TAF8 that interacts with TAF2, are both required for mouse embryonic stem cell survival. Thus, our study demonstrates that distinct TAF8 regions involved in connecting lobe B to lobe C are crucial for TFIID function and consequent ESC survival.

## Introduction

Transcription regulators in eukaryotes can be divided into three functional classes: genespecific transcription regulators, cofactor complexes and the general RNA polymerase transcription machinery. Their collaborative action is necessary to access specific loci in chromatin and allow precise transcription initiation (1). Regulated RNA polymerase II (Pol II) transcription requires a highly concerted, stepwise assembly of transcription factor complexes that form the preinitiation complex (PIC). A functional PIC consists of Pol II and six general transcription factors (GTFs): TFIIA, TFIIB, TFIID, TFIIE, TFIIF and TFIIH (2,3). The evolutionary conserved TFIID complex is the first GTF that binds gene promoters, and along with other GTFs, acts as a platform for PIC formation and consequent transcription initiation (3,4). TFIID is a multisubunit complex of about 1 MDa, composed of the TATA box-binding protein (TBP) and 13 (14 in yeast) TBP-associated factors (TAFs)(5). TFIID not only is essential for the recognition of core promoter sequences and the recruitment of the PIC, but also involved in gene expression via its interactions with cofactors, gene-specific activators and repressors, and chromatin modifications associated with active regions of the genome (4,6,7).

Human TAF8 is a 310 amino acid protein harboring a histone fold domain (HFD) at its N-terminal end, that interacts with the HFD of TAF10, to form a non-canonical histone fold pair arrangement in TFIID (8–10). TAF8 also interacts with TAF2, and TAF2-TAF8-TAF10 subcomplex assembles in the cytoplasm of human cells (10). Biochemical studies revealed that TFIID is assembled in a step-wise manner, first forming a stable 5TAF core complex, consisting of two copies each of TAF5-TAF6-TAF9-TAF4-TAF12. On one hand, this core is bound by the TAF8-TAF10 heterodimer, forming the 7TAF complex, similar to lobe B (11,12), or by TAF8-TAF10-TAF2 complex, forming the 8TAF complex (10,12–14). On the other hand, the TAF5-TAF6-TAF9-TAF4-TAF12 core is bound by TAF11-TAF13 and TAF3-TAF10 HF pairs and TBP to form lobe A (12,14). Importantly, *in vitro* the TAF8-TAF10 HF pair does not interact individually with any other HF TAF pair, but it interacts with the 5TAFcore complex, only if all five TAFs of the core complex are simultaneously present and the entire 5TAF core complex is formed (11). In addition, we demonstrated that the building blocks of mammalian TFIID, such as TAF8-TAF10, TAF6-TAF9 and TBP-TAF1, assemble co-translationally in the cytoplasm, in agreement with the stepwise assembly model of TFIID (15). Early electron microscopy (EM) studies established that endogenous human TFIID resembles a horseshoe composed of three main lobes (16,17). Recent human and yeast *Komagataella phaffii* TFIID cryoEM structures confirmed the three-lobe-structure of TFIID (called lobes A, B and C), and demonstrated evolutionary conservation and high flexibility within TFIID (12,14,18). The high resolution structures of two TFIID domains indicated that i) lobe B contains the HFD domains of TAF8-TAF10 histone fold pair, together with one copy of the 5TAF core (TAF5-TAF6-TAF9-TAF4-TAF12) complex, ii) TAF8 participates in connecting lobe B and C by interacting with the two HEAT repeats of TAF6 and certain regions of TAF1, and iii) in lobe C the C-terminal half of TAF8 interacts with TAF2 (Fig. 1A) (12,14,18).

**Figure 1.**
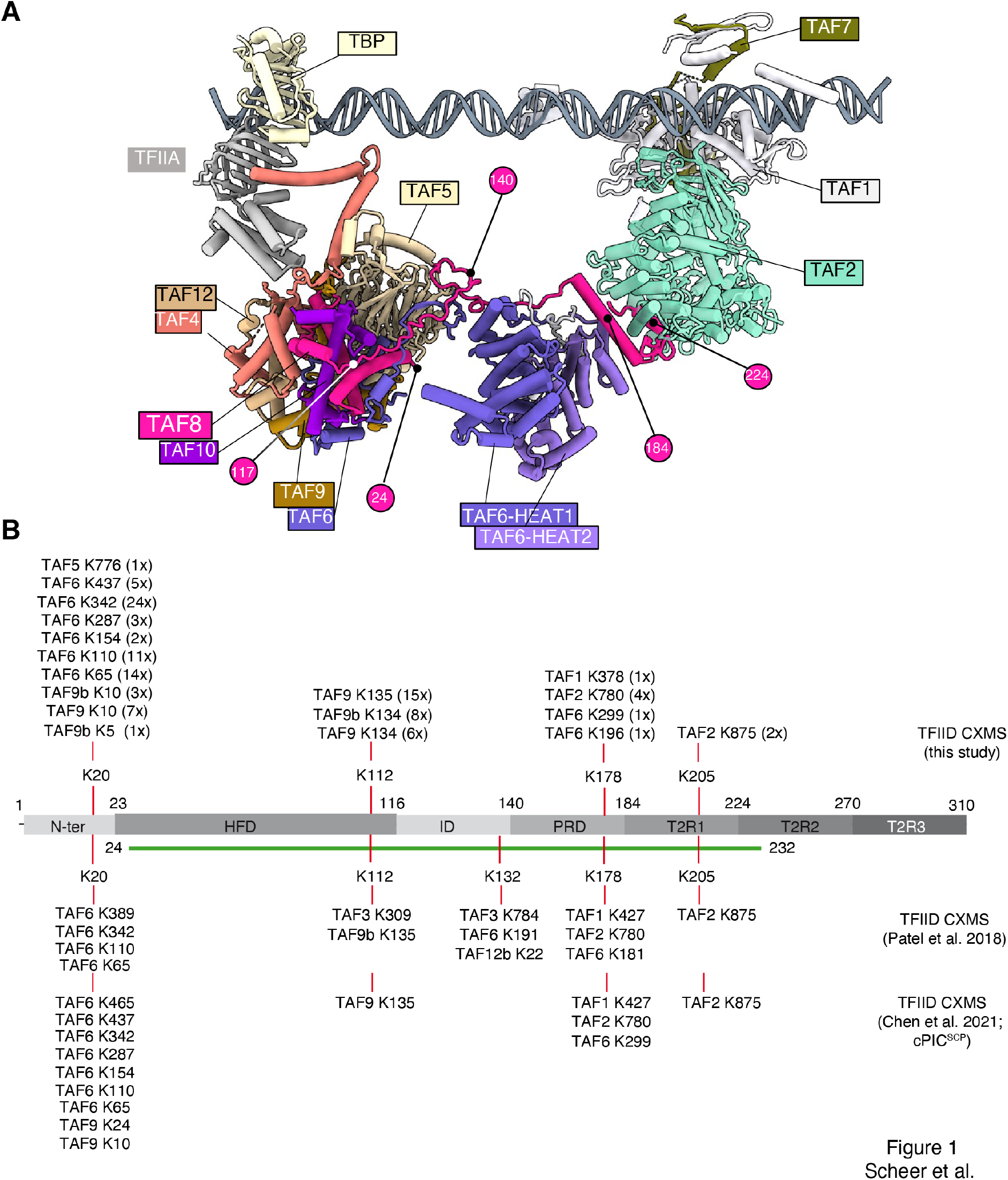
TAF8 interactions in human TFIID. **(A)** The human TFIID cryo-electron microscopy model is shown. The model used in the panel is the super core promoter (SCP)-bound TFIID-TFIIA in post TBP-loading state (12), PDB: 7EGJ). TAF8 is highlighted in rose. TAF8 positions used in our study and visible in the TFIID structure are labelled with rose circles with their corresponding amino acid positions highlighted in white. TBP, TFIIA and TAFs in the TFIID complex are indicated with their corresponding colors. Lobe B is composed of TAF5, 4-12, 6-9 and TAF8-10, and Lobe C of TAF1, TAF2 and TAF7. The HEAT repeats of the two TAF6s connecting the three lobes of TFIID are highlighted. Lobe A is hidden for clarity (lays behind lobe C from this view). (B) TAF8 interlinks (crosslinks between TAF8 and other TFIID subunits) identified by CXMS analyses of our endogenous TFIID complex preparation. All crosslinks were identified with high confidentiality and the number of times a crosslinked peptide was identified is indicated in brackets (Tables S1 and S2). Upper part, CXMS done in this study, lower part, CXMS done by Patel et al. (2018) or Chen et al. (2021). Lysines (K) in TAF8 and the residues to which they crosslink on other TAFs are indicated. Green line indicates the TAF8 path visible in the cryo-EM structure (12).

In mice, germ line knock-out of genes encoding several TFIID subunits (*Taf7, Taf8, Taf10*, and *Tbp*) results in early embryonic lethality around E4.0 (19–22), suggesting that these mammalian TFIID subunits are absolutely essential for early mouse development. Furthermore, conditional deletion of either *Taf8* in mouse embryonic stem cells (ESCs) (23), or *Taf10*, or *Taf4* in embryonic keratinocytes (24,25), or deletion of *Taf10* in embryonic liver (26), or ablation of *Taf7* in CD4^-^CD8^-^ thymocytes (20) compromise the viability of all these mutant cells, suggesting that TFIID subunits play essential roles in transcription in the different tested cellular systems.

Importantly, a homozygous *TAF8* TAF8c.781–1G>A splice site mutation in a human patient causes intellectual disability (23). Patients with this mutation express an unstable TAF8 protein in which the C-terminal 49 wild type amino acids are replaced by a 38 amino acid mutated sequence, caused by the frame shift, leading to partial TFIID dissociation (23). Interestingly, several mutations have also been reported in TAF2 (i.e. T186R, P416H, or W649R), the interaction partner of TAF8, that are also associated with intellectual disability syndrome (27,28).

In the present study, we have investigated the requirement of different TAF8 regions for TFIID assembly, and the importance of some of these TAF8 regions for embryonic stem cell (ESC) viability. We have identified TAF8 regions which are either important for interacting with the 5TAF core complex in lobe B, or for interacting with TAF2 in lobe C, and we show that the 5TAF core-interacting, and that one of the TAF2-interacting TAF8 regions are required for mouse embryonic stem cell survival.

## Results

### Crosslinking mass spectrometry analysis of human TFIID reveals crosslinking hotspots in TAF8

To gain more insights into the structure/function relation of human TAF8, first we carried out a multiple sequence alignment of TAF8 proteins from several eukaryotic species to better understand the potential domain conservation of TAF8 in addition to its HFD (Fig. S1). Following our alignment and based on previous publications (9–11), we have subdivided TAF8 in seven regions: N-ter, HFD, ID (intermediary domain), PRD (proline-rich domain), and T2R1, T2R2 and T2R3 (TAF2-interacting regions 1, 2 and 3) (Fig. 1A, 2A and Fig. S1). Prior to the publication of the cryo-EM structure of hTFIID (14), we performed cross-linking mass spectrometry (CXMS) analyses on TFIID to characterize the architecture of the complex. To this end we purified endogenous TBP-containing complexes (TFIID, SL1 and TFIIIB) by an anti-TBP immunoprecipitation from HeLa cell nuclear extracts, (Fig. S2), crosslinked them with the amine-reactive crosslinker BS3 and analyzed the sample by CXMS. The CXMS analyses of the endogenous TFIID complex identified 65 interlinks (crosslinks between different protein molecules) and 43 intralinks (crosslinks involving the same protein) (Table S1 and Table S2, respectively). We then focused on TAF8 interlinks where TAF8 cross-linked to another TAF peptide and compared our dataset to CXMS data obtained from super core promoter (SCP) bound hTFIID, or SCP bound hTFIID containing PIC (12,14) (Fig. 1B). In all the analyzed endogenous hTFIID complexes TAF8 crosslinked extensively to TAF2, TAF6, TAF9/9b, and with a lesser frequency to TAF1, TAF3, TAF5 and TAF12 with crosslinking hotspots in TAF8 at K:20, K:112, K:132, K:178 and K:205 (Fig. 1B). The K:20 hotspot of TAF8, is absent from the published human TFIID structure (PDB: 6MZM, (14)), and crosslinks to the TAF9/TAF6 HFD, to the TAF6 middle (TAF6M) and TAF6C domains, containing five HEAT repeats (29), and to one position at the C-terminus of TAF5. These crosslinks agree with the potential position of the N-terminal end of TAF8 in the TFIID structure (Fig. 1A and 1B). K:112 at the C-terminus of the HFD of TAF8 crosslinks to a region downstream of the HFD in TAF9/9b and to TAF3. K:132 in ID of TAF8 crosslinks to TAF3, TAF6 and TAF12b. K:178 at the C-terminus of the PRD crosslinks to the non-conserved TAF6 linker between TAF6M and TAF6C, the third HEAT repeat of TAF6, to TAF2 and to the N-terminus region of TAF1. K:205 in T2R1 of TAF8 crosslinks in all the three CXMS analyses to K:875 in TAF2. Surprisingly, between amino acids 205 and 310 of TAF8 (situated in T2R2 and T2R3 regions) no crosslinked peptides were detected in any of the three TFIID preparations (Fig. 1B), suggesting that the C-terminal end of TAF8 does not produce peptides that are amenable to MS analysis, eventually is buried in a non-crosslinkable structure, or is sticking out from the TFIID surface. Note that the visible path of hTAF8 in the cryo-EM structure of the human TFIID is from amino acids 28 to 229 (PDB: 6MZM, (14)) encompassing the HFD, ID, PRD, and T2R1 regions of TAF8 (Fig. 1A). Nevertheless, these crosslinking results in agreement with cryo-EM structures, further indicate that TAF8 plays a triple role in TFIID: i) participates in the assembly of the six HFD-containing 7TAF complex in lobe B via its N-terminus and HFD regions, ii) connects lobe B and C by interacting with the HEAT repeat regions of two TAF6, the TAF1 regions between aa 378-427 and TAF2 via its PRD, and iii) interacts with TAF2 in lobe C via its C-terminal part.

**Figure 2.**
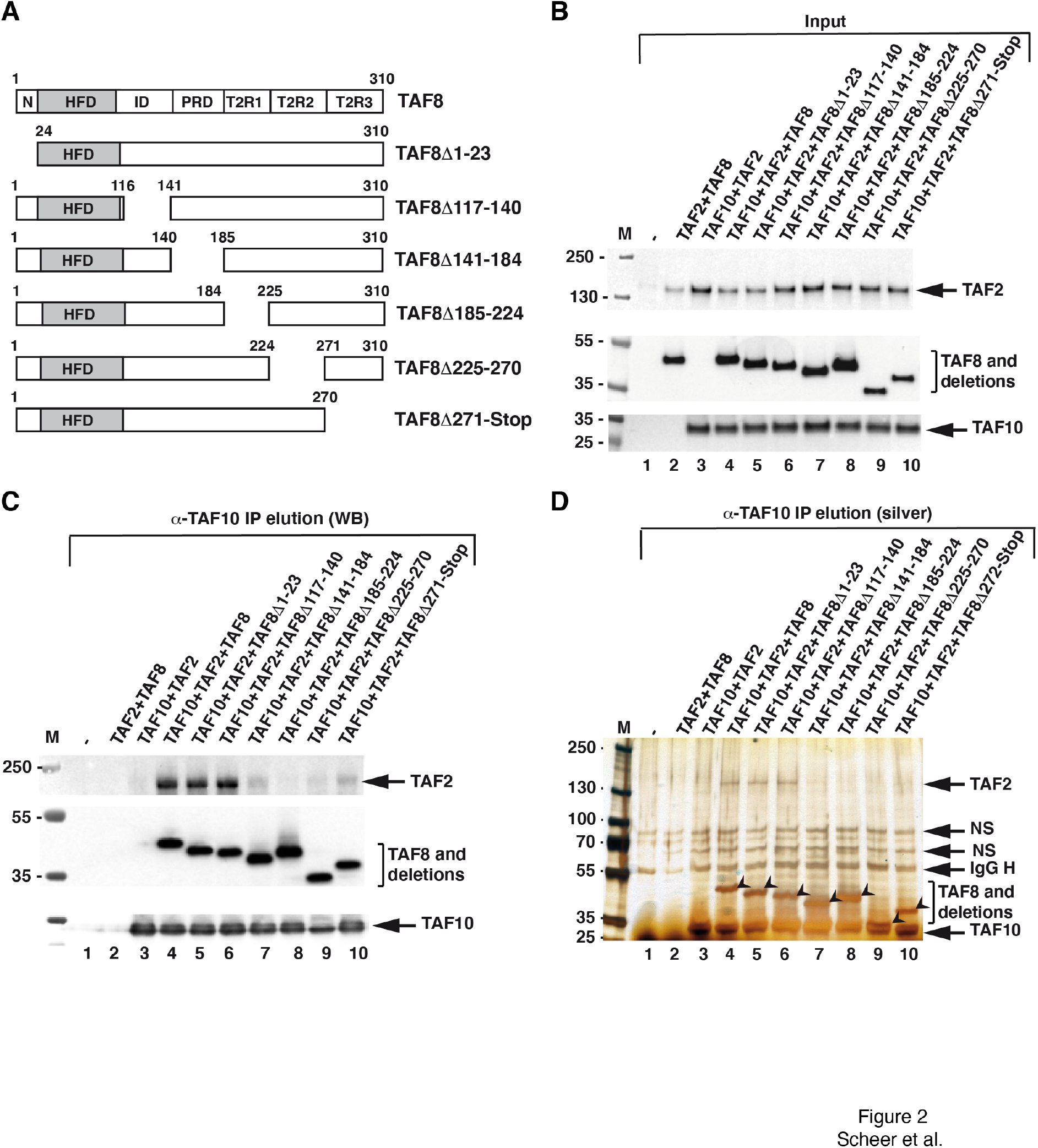
Characterization of TAF8 interactions within the TAF2-TAF8-TAF10 subcomplex. **(A)** Schematic presentation of human TAF8, its domain structure, and the different TAF8 deletions. N: N-terminal domain; HFD: histone fold domain; ID: intermediary domain, PRD: proline-rich domain, and T2R1, T2R2 and T2R3: TAF2-interacting regions 1, 2 and 3. **(B-D)** TAF2 and TAF10 were co-expressed with either WT TAF8 or with TAF8 deletions as indicated above each lane using the baculovirus overexpression system. Input extracts (B) and anti-TAF10 IPs (C) were tested by western blot analyses (WB) with the indicted antibodies. (D) Anti-TAF10 IPs were also tested by silver-staining (silver) of the gels. TAF8 and its deletions are indicated with an arrowhead. On panels (B-D) the molecular weight markers (M) are indicated in kDa. (D) IgG H: heavy chain of the antibody. NS: non-specific band. On panels (B-D) - (lanes 1) indicates Sf9 cell extract without any coinfection.

### Identification of regions of TAF8 that are required either to stably interact with the 5TAF core complex or with TAF2

To further delineate the regions of TAF8 that are important for lobe B and/or lobe C assembly/interactions in TFIID, based on the above determined domain organization (Fig. 1, and Fig. S1) and their cross-linking hotspots, we constructed a series of deletion mutants in TAF8. In these deletion mutants, small regions (about 23-45 amino acids) throughout the whole length of human TAF8 were individually deleted, without deleting the HFD of TAF8 to keep the TAF8-TAF10 HF-interaction interface intact (Fig. 2A). To evaluate the impact of these deletions on the assembly of recombinant 3TAF (TAF10-TAF8-TAF2), 7TAF (TAF5-TAF6-9/9b-TAF4-12-TAF8-TAF10) and 8TAF (7TAF complex+TAF2) complexes, we used the baculovirus expression system.

First, we co-expressed TAF10-TAF2 with wild type (WT) TAF8, or with the six deletion mutants, and carried out TAF10 immunoprecipitations (IPs) (Fig. 2B, 2C and 2D), to test whether TAF8 deletion mutants would abolish interaction(s) of the TAF8-TAF10 HF pair with TAF2. Our anti-TAF10 coimmunoprecipitation (co-IP) experiments indicated that deletion of PRD [amino acids (aa) 141-184], T2R1 (aa 185-224), T2R2 (aa 225-271), or T2R3 (aa 272-Stop) of TAF8, all abrogated TAF8-TAF10 interactions with TAF2 (Fig. 2C and 2D). Deletions of N-terminal (aa 1-23 and ID (aa 117-140) regions of TAF8 did not influence the formation of the 3TAF (TAF10-TAF8-TAF2) complex (Fig. 2C and 2D). In our negative control experiments, when TAF10 was absent from the coinfections, TAF8 or TAF2 did not co-purify by the anti-TAF10 IP, or when TAF8 was absent from the coinfections, TAF10 did not interact with TAF2 (Fig. 2B-2D, lanes 2 and 3).

Next, subunits of the 7TAF complex were co-expressed with either WT TAF8, or with one of the six deletion mutants (Fig. 2A) followed by anti-TAF10 IPs. When analyzing the 7TAF complex formation, we found that only the deletion the ID region of TAF8 (Δ117-140) severely reduced the interactions with subunits of the 5TAF core complex, while the TAF8-TAF10 HF pair could still form (Fig. S2A, Fig. 3A, 3B). All other tested TAF8 deletion mutants, including the delta N-terminus mutant deleting the K:20 crosslinking hotspot, which crosslinked extensively to subunits of the 5TAF complex (Fig. 1B), formed the 7TAF complex (Fig. 3A, 3B). As described previously (11), TAF10 alone could not interact with the 5TAF core complex when TAF8 was absent from the coinfections, further indicating that the formation of the TAF8-TAF10 HF pair is crucial for forming lobe B complex in TFIID (Fig. 3A, 3B).

**Figure 3.**
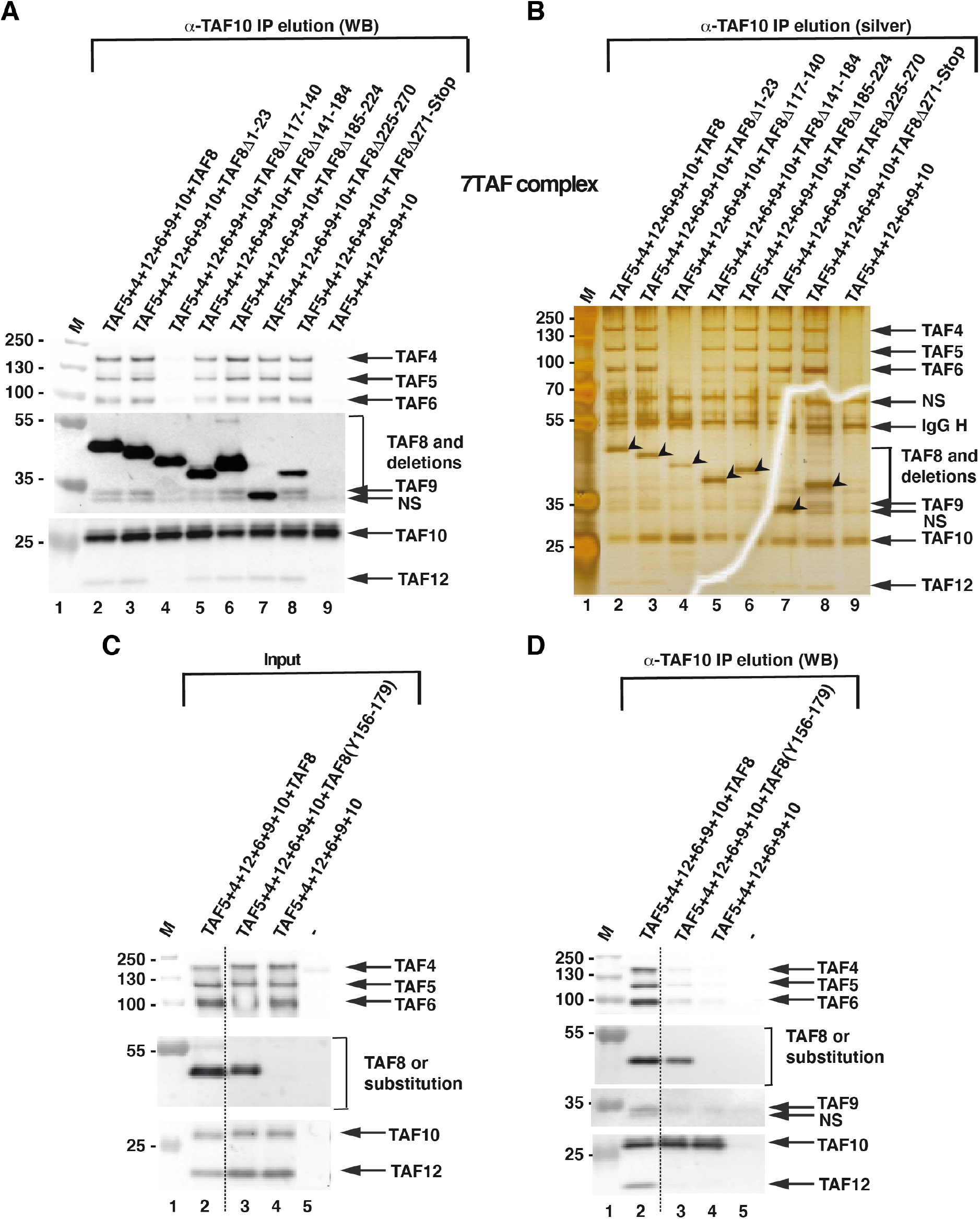
Characterization of TAF8 interactions within the 7TAF complexes. **(A-B)** TAF5-TAF6-TAF9-TAF4-TAF12 and TAF10 were co-expressed with either WT TAF8, or with TAF8 deletions as indicated above each lane using the baculovirus overexpression system. Anti-TAF10 IPs (A) were tested by western blot analyses (WB) with the indicated antibodies, or by silver staining (silver) of the gels (B). Anti-TAF10 IPs (A) were tested by western blot analyses (WB) with the indicted antibodies, or by silver-staining (silver) of the gels (B). On panel (A) the molecular weight markers (M) are indicated in kDa. NS: nonspecific band. (B) The arrow heads indicate TAF8 or the different TAF8 deletions. IgG H: heavy chain of the antibody. NS: non-specific band. The western blot assay tests corresponding to the TAF expressions in the Input extracts for (A-B) are shown in Supp Figure 3. **(C-D)** The substitution of human TAF8 117-140 amino acid sequences with the *S. cerevisiae* Taf8 sequences does not rescue 7TAF complex formation. TAF5-TAF4-TAF12-TAF6-TAF9 and TAF10 were co-expressed with either WT TAF8, or with the TAF8 substitution in which the human 117-140 amino acid (aa) sequences were replaced with the yeast 156-179 aa sequences ([TAF8(Y156-179)] as indicated above each lane) using the baculovirus overexpression system. Input extracts (C) or anti-TAF10 IPs (D) were tested by western blot analyses (WB) with the indicted antibodies. Molecular weight markers (M) are indicated in kDa. -: indicates Sf9 cell extract without any coinfection. Dotted lines indicate where the gels were cut.

To test whether the amino acid sequence of the ID is required for formation of 7TAF complex, we substituted the human TAF8 117-140 amino acid sequences with the *S. cerevisiae* Taf8 sequences from amino acids 156 to 179. TAF10 co-IP results showed that the amino acid substitution in this region could not functionally replace the human sequences, as the 7TAF complex did not form efficiently under these conditions (Fig. 3C, 3D). The results show that the amino acid sequence of the ID domain is required for efficient formation of the 7TAF complex, and that the ID is not functionally conserved between *S. cerevisiae* and humans.

When subunits of the 8TAF complex were co-expressed with either WT TAF8, or with one of the six deletion mutants (Fig. 2A, Fig. S3B) and anti-TAF10 IPs carried out, the results recapitulated the observations made individually with either the 3TAF complex, or with the 7TAF complex (Fig2B-2D, Fig.3). Namely, in the 8TAF experiments the TAF8Δ117-140 mutant abrogated the TAF8-TAF10 interactions with the subunits of the 5TAF core complex (TAF5, 4-12, 6-9), but not with TAF2 (Fig. 4A and 4B), and deletion of PRD (Δ141-184), T2R1 (Δ185-224), T2R2 (Δ225-271) or T2R3 (Δ272-Stop) of TAF8 all severely abrogated the TAF8-10 HF pair interactions with TAF2, but did not influence the interactions with the 5TAF core complex (TAF5, 6-9, 4-12,) (Fig. 4A and 4B).

**Figure 4.**
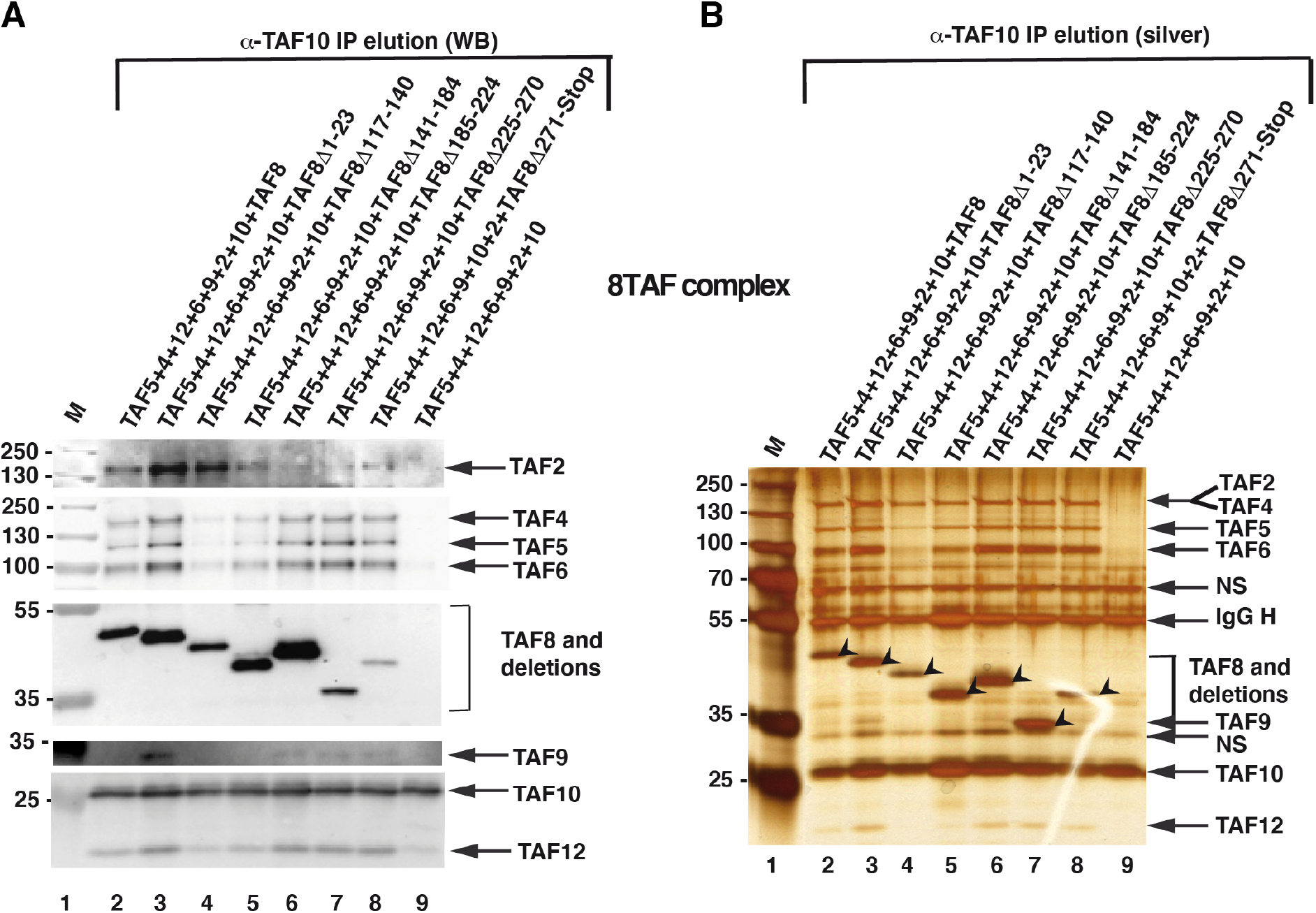
Characterization of TAF8 interactions within the 8TAF complexes. **(A-B)** TAF2-TAF5-TAF6-TAF9-TAF4-TAF12 and TAF10 were co-expressed with either WT TAF8, or with TAF8 deletions, as indicated above each lane, using the baculovirus overexpression system. Anti-TAF10 IPs (A) were tested by western blot analyses (WB) with the indicted antibodies, or by silver-staining (silver) of the gels (B) as described in Figure 3A and B. On panel (A) the molecular weight markers (M) are indicated in kDa. NS: non-specific band. (B) The arrow heads indicate TAF8 or the different TAF8 deletions. Input extracts for the anti-TAF10 IPs are shown in Supp Figure 3.

Thus, together our recombinant *in vitro* TFIID sub-assembly results show that i) deletion of the non-conserved N-terminal region of TAF8, which crosslinks to many core TAF subunits in TFIID (Fig. 1B), is not required for formation of the 7TAF core complex, or to interact with TAF2, ii) the ID region of TAF8 (aa 117-140) is required to interact with the 5TAF core complex and it cannot be replaced by the ID region from *S. cerevisiae* Taf8, and iii) PRD (aa 141-184], T2R1 (aa 185-224), T2R2 (aa 225-271), or T2R3 (aa 272-Stop) of TAF8 are required for stable interaction of TAF8-TAF10 with TAF2.

### Exogenously expressed TAF8 lacking ID, or PRD regions are defective for TFIID assembly in mESCs

To test whether the *in vitro* identified regions of TAF8 play a role in TFIID assembly/function in a cellular context, we created mouse embryonic stem cells (ESCs) expressing Flag-WT TAF8, Flag-TAF8Δ 117-140 (ID) or Flag-TAF8Δ141-184 (PRD) mutants. To generate a Dox-inducible expression system, first we integrated an expression cassette encoding the reverse tetracycline-controlled trans-activator (rtTA; (30)) in E14 ESCs (rtTA-ESC). (Figure S4). Next we integrated in the genome of the selected rtTA-ESC clone (DD1) a series of cassettes in which the *Flag-WT-TAF8, Flag-TAF8Δ117-140* or *Flag-TAF8Δ141-184* cDNAs are under the control of a tetracycline-operator, resulting in the Flag-TAF8WT:R, Flag-TAF8Δ117-140:R or Flag-TAF8Δ141-184:R expressing ESC lines, respectively. The inducibility of Flag-WT-TAF8 protein or its deletions by the tetracycline analogue Dox was verified by Western blot analyses using the anti-Flag antibody (Fig. 5A, 5B and 5C). Barely any Flag-tagged protein could be detected in cells grown in the absence of Dox, whereas after Dox treatment Flag-TAF8 WT, Flag-TAF8Δ117-140 or Flag-TAF8Δ141-184 mutant proteins were clearly detectable (Fig. 5A, 5B and 5C).

**Figure 5.**
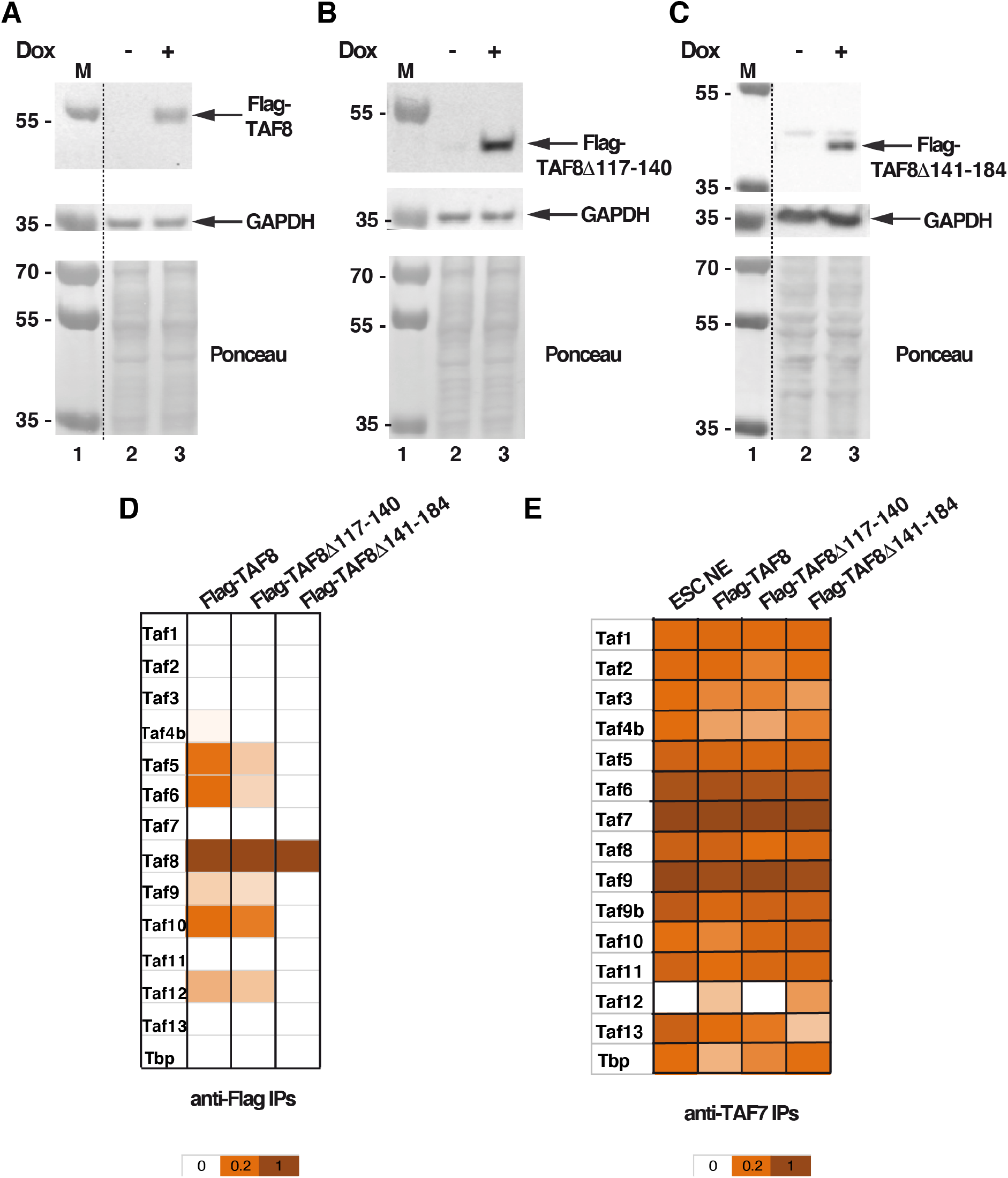
The exogenously expressed Flag-TAF8Δ117-140 and Flag-TAF8Δ141-184 deletions are impaired in TFIID assembly in mESCs. **(A-C)** Stable doxycycline (Dox) inducible E14 mESCs were generated in which WT R:Flag-mTAF8, R:Flag-mTAF8Δ 117-140 or R:Flag-mTAF8Δ141-184 were exogenously expressed. Stable clones were isolated, whole cell extracts (WCE) prepared and tested by western blot assay for the expression of the corresponding Flag-tagged TAF8 proteins (as indicated). Either anti-GAPDH western blot analysis, or Ponceau staining was used as loading control. Molecular weight markers (M) are indicated in kDa. Dotted lines indicate where the gels were cut. **(D, E)** Mass spectrometry analysis of either anti-Flag (D), or anti-TAF7 IPs (E) carried out using WCEs prepared from E14 ESCs expressing either WT Flag-TAF8, Flag-mTAF8Δ117-140, or Flag-mTAF8Δ141-184. Three technical replicates were carried out. Normalized spectral abundance factor (NSAF) values were calculated and normalized to the bait of the IPs (to Flag-tagged TAF8 or its deletions, in D; and to TAF7 in E). The normalized NSAF results are represented as heat maps with the indicated scales.

Next, we tested whether the exogenously expressed Flag-WT-TAF8, Flag-TAF8Δ117-140 or Flag-TAF8Δ141-184 proteins would incorporate in TFIID or corresponding subassemblies and/or would influence endogenous TFIID assembly in mESCs. To this end we prepared whole cell extracts from Dox treated Flag-TAF8 WT:R, Flag-TAF8Δ117-140:R or Flag-TAF8Δ141-184:R expressing cell lines, carried out an anti-Flag or anti-TAF7 IPs and subjected the IP-ed complexes to mass spectrometric analyses. Analyses of the MS data from the Flag IP-ed mESC complexes showed that exogenously overexpressed WT Flag-TAF8 incorporated in a 7TAF-like complex, indicating that the excess of exogenously expressed TAF8 can form a TFIID assembly intermediate, and that this intermediate resembles the 7TAF complex, containing TAF6-9, TAF4b-12, TAF5 as well as TAF8-10 (Fig. 5D). In contrast, the Flag-TAF8Δ117-140 mutant’s incorporation in such a TAF intermediate complex was impaired, as the TAF8Δ117-140 mutant associated with less TAF4B, TAF5 and TAF6. In contrast, Flag-TAF8Δ141-184 did not interact with any endogenous TFIID subunit (Fig. 5D). Importantly, none of the exogenously expressed TAF8 proteins (WT or mutants) impaired endogenous TFIID assembly when analyzed by an anti-TAF7 IP (Fig. 5E). Thus, these results show that the ID or the PRD regions of TAF8 are required for efficient assembly of 7TAF complexes (or TFIID subcomplexes) in mESCs.

### The ID and PRD regions of TAF8 are required for mouse embryonic stem cell survival

To test whether the above identified ID or PRD regions of TAF8 are required for *in vivo* TFIID assembly and consequent mESC survival, we set out to inactivate the endogenous *Taf8* gene using CRISPR/Cas9 genome editing in the ESC lines expressing Flag-TAF8 WT:R, Flag-TAF8Δ117-140:R (ID), or Flag-TAF8Δ141-184:R (PRD) (Figure 6A). Using this strategy in the presence of Dox in the Flag-TAF8 WT:R mESC expressing line, we obtained several viable homozygous *Taf8* deletion ESC lines (Figure 6B), indicating that the CRISPR/Cas9 worked efficiently. By using the same gene editing strategy in the presence of Dox in Flag-TAF8Δ117-140:R or Flag-TAF8Δ141-184:R expressing cell lines, we obtained heterozygous clones in the same proportion as for the Flag-TAF8 WT expressing lines, indicating that the CRISPR/Cas9 worked with comparable efficiency at the *Taf8* genomic locus in these heterozygous mESC lines, as in the Flag-TAF8 WT:R mESC expressing line (Fig. 6B). In contrast, we could not isolate any homozygous knockout clone when inactivating the *Taf8* locus in the Flag-TAF8Δ117-140:R or in the Flag-TAF8Δ141-184:R mESC lines, suggesting that the deleted ID or PRD regions of TAF8 have essential roles in endogenous TFIID assembly and/or function, and consequently for mESC survival, and that these functions cannot be compensated by other TAFs or TAF assemblies.

**Figure 6.**
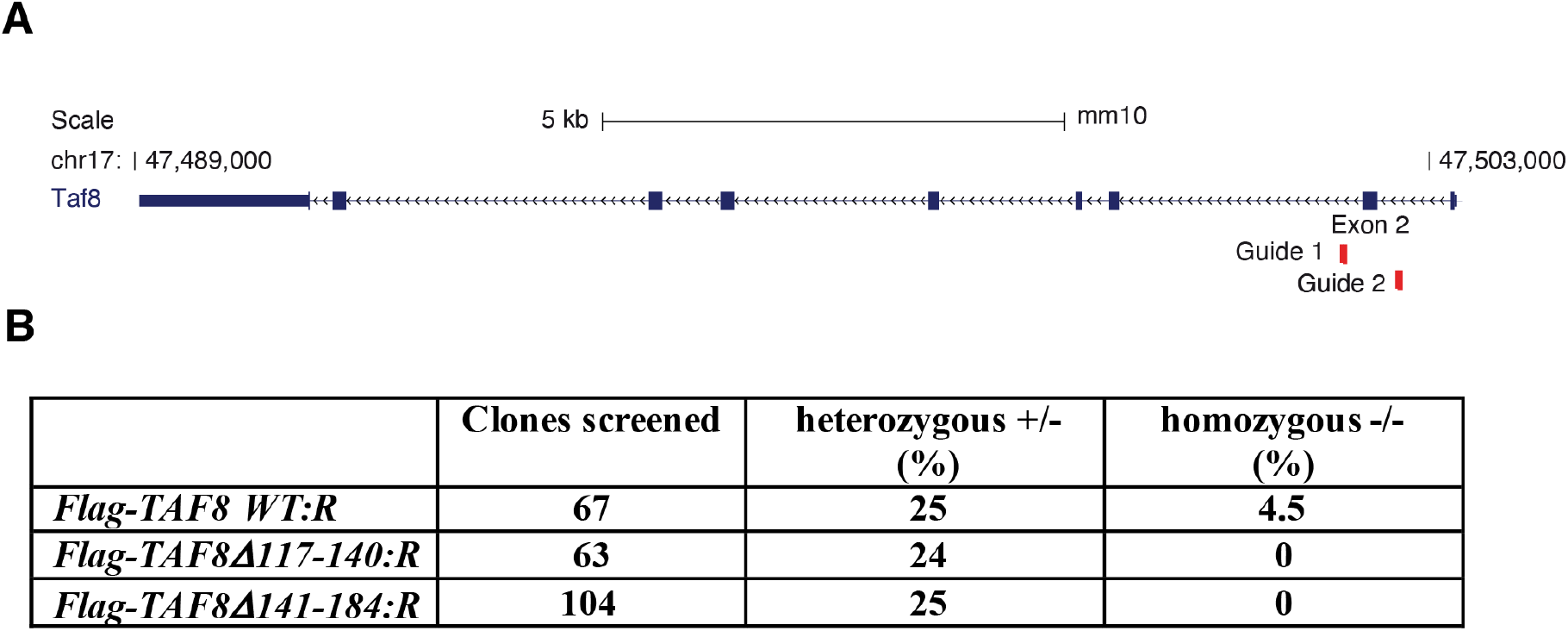
The expression of Flag-TAF8Δ117-140, or Flag-TAF8Δ141-184 do not rescue the knockout of the endogenous *Taf8* gene in mESCs. **(A)** Schematic representation of the mouse *Taf8* locus, with exons and introns and with the direction of transcription, are shown. The genomic positions of the mouse *Taf8* locus on chromosome 17 (chr17) are indicated. The positions of the two gRNAs, surrounding exon 2 used to knockout the endogenous *Taf8* locus, are indicated. **(B)** Table showing the numbers of individual viable ESC clones (R:Flag-mTAF8, R:Flag-mTAF8Δ 117-140 or R:Flag-mTAF8Δ141-184) screened following the CRISPR/Cas9 KO of the endogenous *Taf8* gene. Percentages of the obtained heterozygous (+/-) and homozygous ESC clones (-/-) are indicated.

## Discussion

Characterization of the structure-function relationship of multisubunit transcription complexes is crucial to understand gene regulation. Whereas three-dimensional models of fully assembled multiprotein complexes derived mainly from single-particle cryo-electron microscopy provide a wealth of information on the architecture of such complexes, the requirement of the precise subunit-subunit and domain-domain interactions to maintain the architecture and function of these gene regulatory complexes, such as TFIID remain largely elusive.

Based on several functional and structural studies (9,10,14,19,31,32), as well as on two independent CXMS experiments carried out with endogenous TFIID (this study and (12,14)), we have subdivided TAF8 into seven regions (Fig. 2A). Out of these seven regions the HFD of TAF8 has already been characterized (10), thus we have investigated how the deletion of the remaining six regions of TAF8 influence the *in vitro* assembly of TAF8-containing TFIID building blocks. We identified three types of regions: i) the N-terminus of TAF8 that did not disrupt the tested interactions with either the 5TAFcore complex or with TAF2, ii) the ID region that is required for interactions with the 5TAF core complex, but does not influence the TAF2 interactions, and iii) four successive TAF2 regions which are all required for normal TAF8-TAF2 interactions, but do not influence the interactions with the 5TAF core complex (Figs 2 and 3). Our TAF8-TAF2 interaction results agree with previous surface plasmon resonance (SPR) experiments using immobilized full-length TAF2 as ligand and maltose binding protein (MBP) fusions of TAF8 fragments (10). Thus, our deletion analysis confirms that TAF8 plays a triple role in TFIID: The N-terminus, the HFD and ID regions of TAF8 participate in the assembly of the six HFD-containing 7TAF complex in lobe B. Importantly, when the ID region of human TAF8 was replaced with the non-conserved amino-acids from yeast TAF8, this replacement did not restore the binding of the TAF8-TAF10 HF pair to the 5TAF core complex, further substantiating that this region and its amino sequence is important for forming the 7TAF complex in lobe B. It is possible that residues within the ID make specific contacts with TAF8 or TAF6 or other components of the 5TAF complex that are required for proper interaction of TAF8-TAF10 with the 5TAF complex. In spite of the fact that TAF8 PRD crosslinks to the TAF6 HEAT repeat regions, deletion of TAF8 PRD (aa 141-184) did not influence the formation of the 7TAF complex, suggesting that the TAF8 PRD is involved in the lobe B-C connecting function of TAF8, but not important for lobe B formation. In the cryo-EM structures of TFIID the lobe B-C connecting function of TAF8 was proposed to involve the unstructured PRD of TAF8. Interestingly, our deletion mutant analyses could not separate the lobe B-C connecting function of TAF8 from its TAF2 interacting function. Instead we found that the deletion of either the PRD, or each of the three TAF2 interaction regions (T2R1-T2R3 mutants) individually, all abrogated the interaction of TAF8 with TAF2. It is conceivable that the simple shortening of the 170 amino acid long C-terminal part of TAF8 causes rupture of the TAF8-TAF2 interactions. However, our present deletion experiments, together with the above mentioned SPR experiments using short TAF8 fragments (10), would rather suggest that the entire 170 amino acid TAF8 tail is needed to interact with TAF2. These interactions could happen through several TAF8-TAF2 contact surfaces, which would be all synergistically required for these interactions. In the cryo-EM structures of TFIID the structured TAF8 path stops at amino acid 229 (Fig. 1A) (14), and similarly the unresolved regions of TAF2 could also be involved in TAF8 interactions. Nevertheless, no TAF8-TAF2 crosslinks have been detected in endogenous TFIID complexes either in T2R2 or T2R3 regions. In this respect it is interesting to note that the deletion of the T2R2 or T2R3 regions of TAF8, both abrogate the TAF2 interactions, suggesting that these regions are still crucial for TAF2 interactions. Intriguingly, the patient with intellectual disability expresses an unstable TAF8 protein in which the C-terminal last 49 wild type amino acids of TAF8 are replaced by a 38 amino acid mutated sequence, caused by the frame shift, leading to potential TFIID dissociation (23). In addition, the reported intellectual disability causing TAF2 mutations (T186R, P416H, or W649R) may also fall in these unmapped TAF2-TAF8 interaction regions (27,28). It is thus conceivable that an important regulatory surveillance mechanism exists in cells to control TAF8-TAF2 interactions and through these interactions stable holo-TFIID assembly. Nevertheless, pluripotent ESCs with highly active Pol II transcription seem to require fully assembled and functional holo-TFIID as deletion of *Tbp, Taf7, Taf8*, and *Taf10* cause mESC lethality (19–22). Our results show that when the ability of TAF8 to interact with the 5TAF core complex in lobe B (TAF8ΔID), or with TAF2 in lobe C (TAF8ΔPRD) are impaired, likely causing defective TFIID assembly, the ESCs cannot survive, further suggesting that ESCs require fully assembled holo-TFIID for function. Future experiments will decide whether such deletions at later stages of ESC differentiation (i.e. embryonic body or neuronal differentiation) would also cause cell death. Here we have identified regions of TAF8 that are required for 1) anchoring TAF8-TAF10 to the core 5TAF complex and 2) for connecting the 7TAF complex in Lobe B to Lobe C. TAF8 plays a critical role in the genesis of TFIID and mutations in TAF8 can lead to human diseases such as ID. As such, defining the role of TAF8 regions in TFIID assembly and function provides key information about how TFIID assembly occurs and how mutations in particular domain can lead to human diseases.

Altogether, we show the significance of precise interactions of TAF8 with other TAFs, establishing TAF8 as a functional bridge between lobes B and C in TFIID. Moreover, our experiments demonstrate a functional role of these interactions in TFIID, required for proper functioning of mESCs.

## Experimental Procedures

### Plasmids

The baculovirus expression vector for FLAG-tagged human (h) TAF8 WT was PCR amplified from the pPBAC MultiBac vector, expressing TAF8 and TAF10 together, described in (10), and the cDNA was inserted in the pVL1393 baculovirus expression vector digested with Bam HI and Eco RI together with a primer coding for a Flag-tag on the 5’ end of the hTAF8 cDNA. The TAF8 deletion series was generated by site directed mutagenesis using pVL1393-Flag-TAF8 WT vector as a template (see Fig. 2A). The deleted Flag-TAF8 cassettes were inserted into the pVL1393 baculovirus expression vector. The constructs were verified by sequencing. All the other baculovirus expression vectors have been described previously (11,33).

Mouse (m) WT Flag-TAF8 cDNA and its deletions, corresponding to the hTAF8 deletions (called Flag-mTAF8Δ117-140 or Flag-mTAF8Δ141-184), were cloned into the pUHD10-3 vector (34) in which the expression of WT Flag-mTAF8, Flag-mTAF8Δ117-140, or Flag-mTAF8Δ141-184 are under the control of tet-operator.

The pUES-3 plasmid expressing the two *Taf8* guide (g) RNAs and co-expressing high-fidelity Cas9 (35) fused to EGFP (Cas9-HF-EGFP) were generated by Golden Gate cloning (36). The sequences of the gRNAs are shown in Table S3. The pUES-3 plasmid was verified by sequencing.

### Antibodies

Mouse monoclonal (mAb) and rabbit polyclonal (pAb) antibodies raised against the following proteins have been described previously, or were purchased commercially: anti-TBP (mAb 3G3) (37) and (mAb 2C1) (38), anti-TAF2 (pAb 3083) (10), anti-TAF4 (mAb 32TA2B9) (22), anti-TAF5 (mAb 1TA1C2) (39), anti-TAF6 (mAb 25TA2G7) (39), anti-TAF7 (mAb 19TA) (40) and (pAb 3475) (41), anti-TAF8 (pAb 3478) (41), anti-TAF10 (mAb 6TA2B11) (42) and TAF12 (22TA) (43), anti-VP16 mAbs 2GV4, 5GV7 and 5GV2 (44), anti-γ-Tubulin (Sigma Aldrich T6557), and anti-FLAG [M2 antibody (F3165 Sigma-Aldrich)].

### Mouse ESC culture conditions

Mouse E14 ESCs were cultured on plates coated with 0.1% gelatin solution in 1x PBS (Dutcher, Cat# P06-20410) using DMEM (4.5 g/l glucose) with 2 mM Glutamax-I medium supplemented with 15% foetal calf serum (FCS) (ThermoFisher Scientific, Cat# 10270-106), 0.1% β-mercaptoethanol (ThermoFisher Scientific, Cat# 31350-010), 40 μg/ml gentamicin (KALYS, Cat# G0124-25), 0.1 mM non-essential amino acids (ThermoFisher Scientific, Cat# 11140-035), 1.500 U/ml leukaemia inhibitory factor (LIF) (home-made), and 3 μM CHIR99021 (Axon Medchem, Cat# 1386) together with 1 μM PD0325901 (Axon Medchem, Cat# 1408) were added freshly to the medium. Cells are grown at 37°C with 5% CO2. For cell collection and/amplification ESCs were trypsinized for 2 to 3 minutes with trypsin-EDTA (Invitrogen, Cat#25200-072) and the digestion was stopped by the addition of pre-warmed 15% FCS. Where indicated doxycycline was added to the medium at a final concentration of 1μg/ml (Sigma Cat#D9891).

### Generation of conditional *Taf8* mutant mouse embryonic stem cells

To obtain an mESC line stably expressing the reverse tetracycline activator rtTA, first E14 mESCs were transfected with the pDG1-rtTA plasmid (Aat II linearized), together with 1/10 of pKJ1B plasmid (for neomycin G-418 selection; Hind III linearized) as described previously (30,45), using Lipofectamine2000 following manufacturer’s instruction (ThermoFisher Scientific, Cat# 11668-019). To select E14 mESCs clones stably expressing rTA, first genomic PCR was used with primers QY74 and WK31 (Table S4). Next, RNA was prepared with NucleoSpin^R^ RNA mini isolation kit (MACHEREY-NAGEL, GmbH&Co.KG, Cat#740955), and RT-qPCR was carried out using HA273 and HA274 primers for *rtTA* amplification, HA817 and HA818 primers for *Rplp0* housekeeping gene amplification (Table S4). The expression of the rtTA protein was verified by western blot analyses using anti-VP16 mAbs 2GV4 5GV7 and 5GV2 (44).

Next the rtTA expressing ESC line was transfected using Lipofectamine2000 (ThermoFisher Scientific, Cat# 11668-019) following manufacturer’s instructions, with pUHD-Flag-mTAF8, pUHD-Flag-mTAF8Δ117-140, or pUHD-Flag-mTAF8 Δ141-184 plasmids (each Hind III linearized), in which the expression of wt mTAF8, and the two mTAF8 deletion mutants are under the control of tet-operator, together with 1/10 of pGK Hygro plasmid (for hygromycin selection; PvuII linearized). Hygromycin (Sigma Cat#HO654) selection with a 250 μg/ml was started two days after transfection (45). Single cell derived colonies were isolated 8 days after hygromycin selection, and amplifed in 3 cm wells. After inducing the expression of the different *Taf8* transgenes for 3 days with 1 μg/ml doxycycline (Dox, D9891 Sigma-Aldrich) positive cell clones were selected by western blot assays using an anti-Flag antibody, or purified anti-TAF8 (pAb3478) (Figure 4A-C).

### Knocking out the endogenous Taf8 locus

R:Flag-mTAF8, R:Flag-mTAF8 Δ117-140, and R:Flag-mTAF8 Δ141-184 E14 derived cells at a confluency of 70-80% (2×10^6^ cells on a 10 cm petri dish) were transfected with 24 μg of a pUES-3 plasmid containing the two *Taf8 gRNAs* (Figure 6A and Table S3) as well as expressing the Cas9-GFP fusion protein using Lipofectamine 2000 (ThermoFisher Scientific, Cat# 11668-019). Dox was omitted during transfection and was added 6 hours after transfection. Two days later, the transfected R:Flag-mTAF8, R:Flag-mTAF8 Δ117-140, and R:Flag-mTAF8 Δ141-184 cells were selected for the expression of the Cas9-GFP fusion protein by fluorescence activated cell sorting (FACS). Five 96-well plates were seeded with one GFP positive cell per well using the BD Biosciences FACSAria Fusion apparatus equipped with Biosafety Cabinet, and single cells were cultured for 8 days, media was changed every two days. Mouse ESC clones originating from a single transfected ESC were collected, transferred to individual wells of a 48-well plates, amplified in the presence of dox and 1/5th of the cells were used for PCR genotyping. The genomic DNA was extracted, and using Phire direct PCR kit (Thermo Scientific, Cat# F-1265) following manufacturer’s instruction. Primer sequences for PCR are shown in Table S4.

### RT-qPCR

For RT-qPCR, the isolated RNA samples were reverse transcribed to cDNA using superscript II (Transcriptase inverse SuperScript™ II, Invitrogen™ Cat#18064022) following manufacturer’s instruction. Then the cDNA samples were amplified using LightCycler^®^ 480 SYBR^®^ Green 2x PCR Master Mix I (Roche, Cat# 04887352001) and 0.6 μM of forward and reverse primers with a LightCycler^®^ 480 (Roche). The primer pairs used for different RT-qPCR reactions are listed in Table 2. For the assessment of mRNA levels, the obtained threshold-values were used to calculate the relative gene expression using the 2-ΔΔCT method and considering the individual primer pair efficiencies (46).

### Whole cell extract preparations

The required number of cells were trypsinized, transferred to 1.5 ml Eppendorf tubes, centrifuged at 600 g 4°C for 2 min, and washed once with 1ml 1x PBS. Pellets were resuspended in 1 packed cell volume (PCV) extraction buffer (600 mM KCl, 50 mM Tris-HCl pH 7.9, 25% glycerol, 0.2 mM EDTA, 0.5 mM DTT, 5 mM MgCl_2_, 0.5% NP40, 1x protease inhibitor cocktail), incubated 5-10 min on ice and mixed with 3 PCV of “IP 0buffer” (25 mM Tris-HCl pH 7.9, 5% glycerol, 1 mM DTT, 5 mM MgCl_2_, 0.1% NP40, 1x protease inhibitor cocktail). After a further 10 min incubation on ice, tubes were centrifuged at 14000 g 4°C for 10 min. The supernatant fractions, called whole cell extracts (WCEs) were stored at minus 80°C.

### Recombinant protein production

Recombinant baculoviruses were generated as described (10);Demeny, 2007 #3366} and used for protein complex production in *Sf9* insect cell culture. Infected insect cells were harvested 48 h post cell infection by centrifugation and stored at −80 °C until further use. Pellets of baculovirus-infected *Sf9* insect cells co-expressing the different TAFs and the indicated TAF8 proteins were resuspended in lysis buffer (400 mM KCl, 50 mM Tris-HCl pH 7.9, 10% glycerol, 0.2 mM EDTA, 0.5 mM DTT, containing 1x protease inhibitor cocktail) and crude extracts were prepared by three rounds of freeze-thawing, and cleared by centrifugation.

### Immunoprecipitation experiments

Protein-G (Protein G Sepharose^®^ 4 Fast, cat# GE17-0618-01) or Protein-A (ProteinA-Sepharose^®^ 4, cat#P9424 Millipore) beads were washed twice with 1x PBS and twice with IP100 buffer (25 mM Tris-HCl 7.9, 5 mM MgCl_2_, 10% glycerol, 0,1% NP40, 100 mM KCl, 2 mM DTT and 1x protein inhibitor cocktail). Starting input protein extracts were ESC nuclear extract: 0.5-2 mg, ESC cytoplasmic extracts: 1.5-6 mg, or ESC WCE: 0.5-2 mg, Baculo recombinant WCE 2-5 mg). If needed, protein extracts were diluted with 0 buffer (25 mM Tris-HCl pH 7.5, 5 mM MgCl_2_, 10% glycerol, 0.1% NP40, 1 mM DTT and 1x protease inhibitor cocktail) to reach 100 mM KCl in the extracts. Protein inputs were then pre-cleared with 1/10 volume of 100% protein A or G beads for 1 hours at 4°C with overhead agitation. Beads were coupled to the different antibodies (as indicated in the figure legends). Approximately, 1 mg of indicated antibody per ml of protein A or G bead was bound. Beads were incubated for 1h at room temperature with agitation, unbound antibody were removed by washing the beads twice with IP500 buffer (0 buffer containing 500 mM KCl) and twice with IP100 buffer (0 buffer containing 100 mM KCl) before addition of the pre-cleared protein extracts, and further incubated overnight at 4°C with overhead agitation. The following day the beads were collected, and subjected to two rounds of washing for 10 minutes each with 10 volumes of IP500 buffer, followed by 2 x IP100 buffer washes. Proteins IP-ed with an anti-TBP or anti-TAF10 mAb, were eluted by adding 1 bead volume of 2 mg/ml competing epitope peptide for 4 hours, repeated again for 2 hours (42). For anti-Flag and anti-TAF7 co-IPs, proteins were eluted with 0.1 M pH 2.8 glycine, then neutralized with 1.5 M Tris pH 8.8. Eluted proteins were either separated on an 4-12% SDS-PAGE gel, along with the input extract, and were probed with antibodies as indicated in the different figures, silver-stained, or analyzed by mass spectrometry.

### Chemical crosslinking mass spectrometry

100 μl of eluted TBP-containing complexes were incubated with 5 mM BS3 in 900 μl total volume of crosslinking buffer (20 mM HEPES, pH 7.9, 0.2 mM EDTA, 2 mM MgCl_2_, 3% Trehalose, 300 mM KCl, 0.01% NP-40, 1 mM TCEP, 0.5 mM PMSF) at room temperature for 2 h. The reaction was quenched by the addition of 2 μl 1 M ammonium bicarbonate. The crosslinked proteins were TCA precipitated and treated as described (47). Mass spectrometry and identification of BS3 crosslinked peptides was performed as described previously (47). The CXMS data have been deposited deposited to the ProteomeXchange Consortium via the PRIDE (48) partner repository with the dataset identifier PXD026575.

### LC MS/MS Mass spectrometry analyses

Protein mixtures were precipitated with TCA (Sigma Aldrich, Cat# T0699) overnight at 4°C. Samples were then centrifuged at 14000 g for 30 minutes at 4°C. Pellets were washed twice with 1 ml cold acetone and centrifuged at 14000 g for 10 minutes at 4°C. Washed pellet were then urea-denatured with 8 M urea (Sigma Aldrich, Cat# U0631) in Tris-HCl 0.1 mM, reduced with 5 mM TCEP for 30 minutes, and then alkylated with 10 mM iodoacetamide (Sigma Aldrich, Cat# I1149) for 30 minutes in the dark. Both reduction and alkylation were performed at room temperature and under agitation (850 rpm). Double digestion was performed with endoproteinase Lys-C (Wako, Cat# 125-05061) at a ratio 1/100 (enzyme/proteins) in 8 M urea for 4 hours, followed by an overnight modified trypsin digestion (Promega, CAT# V5113) at a ratio 1/100 (enzyme/proteins) in 2 M urea for 12 hours.

Samples were analyzed using an Ultimate 3000 nano-RSLC (ThermoFisher Scientific, San Jose California) coupled in line with a LTQ-Orbitrap ELITE mass spectrometer via a nanoelectrospray ionization source (ThermoFisher Scientific, San Jose California). Peptide mixtures were loaded on a C18 Acclaim PepMap100 trap-column (75 μm ID x 2 cm, 3 μm, 100Å, ThermoFisher Scientific) for 3.5 minutes at 5 μL/min with 2% ACN (Sigma Aldrich, Cat# 1207802), 0.1% formic acid (Sigma Aldrich, Cat# 94318) in water and then separated on a C18 Accucore nano-column (75 μm ID x 50 cm, 2.6 μm, 150Å, ThermoFisher Scientific) with a 90 minutes linear gradient from 5% to 35% buffer B (A: 0.1% FA in water/ B: 99% ACN, 0.1% FA in water), then a 20 minutes linear gradient from 35% to 80% buffer B, followed with 5 min at 99% B and 5 minutes of regeneration at 5% B. The total duration was set to 120 minutes at a flow rate of 200 nL/min. The oven temperature was kept constant at 38°C. The mass spectrometer was operated in positive ionization mode, in data-dependent mode with survey scans from m/z 350-1500 acquired in the Orbitrap at a resolution of 120,000 at m/z 400. The 20 most intense peaks (TOP20) from survey scans were selected for further fragmentation in the Linear Ion Trap with an isolation window of 2.0 Da and were fragmented by CID with normalized collision energy of 35%. Unassigned and single charged states were rejected. The Ion Target Value for the survey scans (in the Orbitrap) and the MS2 mode (in the Linear Ion Trap) were set to 1E6 and 5E3 respectively and the maximum injection time was set to 100 ms for both scan modes. Dynamic exclusion was used. Exclusion duration was set to 20 s, repeat count was set to 1 and exclusion mass width was ± 10 ppm.

#### Data analysis

Proteins were identified by database searching using SequestHT (ThermoFisher Scientific) with Proteome Discoverer 2.4 software (PD2.4, ThermoFisher Scientific) on *Mus musculus* database (Swissprot, non-reviewed, release 2019_08_07, 55121 entries). Precursor and fragment mass tolerances were set at 7 ppm and 0.6 Da respectively, and up to 2 missed cleavages were allowed. Oxidation was set as variable modification, and Carbamidomethylation as fixed modification. Peptides were filtered with a false discovery rate (FDR) at 1%, rank 1 and proteins were identified with 1 unique peptide. For the Label-Free Quantification, the protein abundancies were calculated from the average of the peptide abundancies using the TOP N (where N = 3, the 3 most intense peptides for each protein), and only the unique peptide were used for the quantification. Data were analyzed by calculation of NSAF (41,49). The mass spectrometry proteomics data have been deposited to the ProteomeXchange Consortium via the PRIDE (48) partner repository with the dataset identifier PXD026575.

## Supporting information

Supplemental Tables 3 and 4, and four Supplemental Figures

## Acknowledgments

We are grateful to the IGBMC cell culture and flow cytometry services for assistance. We thank D. Schmit for starting the initial experiments on this project, F. El-Saafin, and the Tora lab members for helpful discussions, A. Bernardini for critically reading the manuscript and for help with generating Figure 1A, N. Jung and B. Reina San Martin for help with the CRIPR/Cas9 experiments.

## Funding and additional information

This study was supported by grants from Agence Nationale de la Recherche (ANR): ANR-19-CE11-0003-02, ANR-PRCI-19-CE12-0029-01, ANR-20-CE12-0017-03, NIH 5R01GM131626-02 and NSF (Award Number:1933344) grants (to LT); NIH R01 GM110064 and GM136974 (to J.R) and supported by funds from CNRS, INSERM, Strasbourg University, and Investissements d’Avenir grants (ANR-10-IDEX-0002-02 and ANR-10-LABX-0030-INRT). The content is solely the responsibility of the authors and does not necessarily represent the official views of the National Institutes of Health or other granting agencies.

## Author Contributions

ES designed and performed most experiments. JL performed the XLMS experiments and their analyses. J-MG generated plasmids and IK-C carried out baculovirus over-expressions. FR carried out the TAF8 IP-related proteomic analyses, KG and IB provided expertise with the TAF complex assemblies. LT and JR supervised the project, and designed the experiments. JR, JL, IB and LT wrote the manuscript.

