## Supplemental Tables 3 and 4, and four Supplemental Figures for "TAF8 regions important for TFIID lobe B assembly, or for TAF2 interactions, are required for embryonic stem cell survival"

**Supporting information:**

**Supporting Tables 3 and 4, and Supporting Figures with legends**

### TAF8 regions important for TFIID lobe B assembly, or for TAF2 interactions, are required for embryonic stem cell survival

Elisabeth Scheer[1,2,3,4], Jie Luo[6], Frank Ruffenach[1,2,3,4], Jean-Marie Garnier[1,2,3,4], Isabelle Kolb-Cheynel[1,2,3,4], Kapil Gupta[5], Imre Berger[5], Jeff Ranish[6], and László Tora[1,2,3,4,*]

[1]Institut de Génétique et de Biologie Moléculaire et Cellulaire, 67404, Illkirch, France

[2]Centre National de la Recherche Scientifique, UMR7104, 67404, Illkirch, France

[3]Institut National de la Santé et de la Recherche Médicale, U964, 67404, Illkirch, France

[4]Université de Strasbourg, 67404, Illkirch, France

[5]School of Biochemistry and Bristol Research Centre for Synthetic Biology BrisSynBio, University of Bristol, Bristol BS8 1TD, UK

[6]Institute for Systems Biology (ISB), Seattle, WA 98109, USA

Key words: TAF8, TFIID, structure, function, TATA binding protein (TBP), TBP-associated factors (TAFs), TAF complexes, embryonic stem cells (ESCs), CRISPR/Cas9, knock out, viability, baculovirus over expression,

Running Title: TAF8 structure/function

**Table S3 List of sgRNAs used to generate *mTAF8*$^{-/-}$ mouse ES E14 cell lines**

| Name | Sequence | PAM | StrandScore (OFF/ON) |
|---|---|---|---|
| mTaf8-1 5' region | 5'- CTGGAGCAAAGTTACAGGAG-3' | CGG | 59/67 |
| mTaf8-2 3' region | 5'- TGCAGACGACTGAAATGACA-3 | AGG | 58/66 |

**Table S4 Primers used for PCR:**

| Primer name | Amplification target gene | Sequence |
|---|---|---|
| PCR primer QY74 Forward | *rtTA* | 5'-CACGCTTCAAAAGCGCACGT-3' |
| PCR primer WK31 Reverse | *rtTA* | 5'-CAATACAGTGTAGGCTGCTC-3' |
| RT qPCR HA273 Forward | *rtTA* | 5'-CCAGCCTTCTTATTCGGCCT-3' |
| RT qPCR HA274 Reverse | *rtTA* | 5'-CCCTCGATCCTAGACCCGTA-3' |
| RT qPCR HA817 Forward | *RPLPO* | 5'-TTGTGAGTGATGTGCAGCTG-3' |
| RT qPCR HA818 Reverse | *RPLPO* | 5'-GGAGATGTTCAGCATGTTC-3' |
| PCR primer 1 Forward | *mTaf8 intron* | 5'- CTCTCCTCTGCATCCTGTGGTGGTACC -3' |
| PCR primer 4 Reverse | *mTAF8 exon* | 5'- ATGGTAGTTATCAGCAGGGTTAGTG -3' |
| PCR primer 6 Reverse | *mTAF8 intron* | 5'- AGAGATTGATTGATTGATTGATTGATTAGG - 3' |

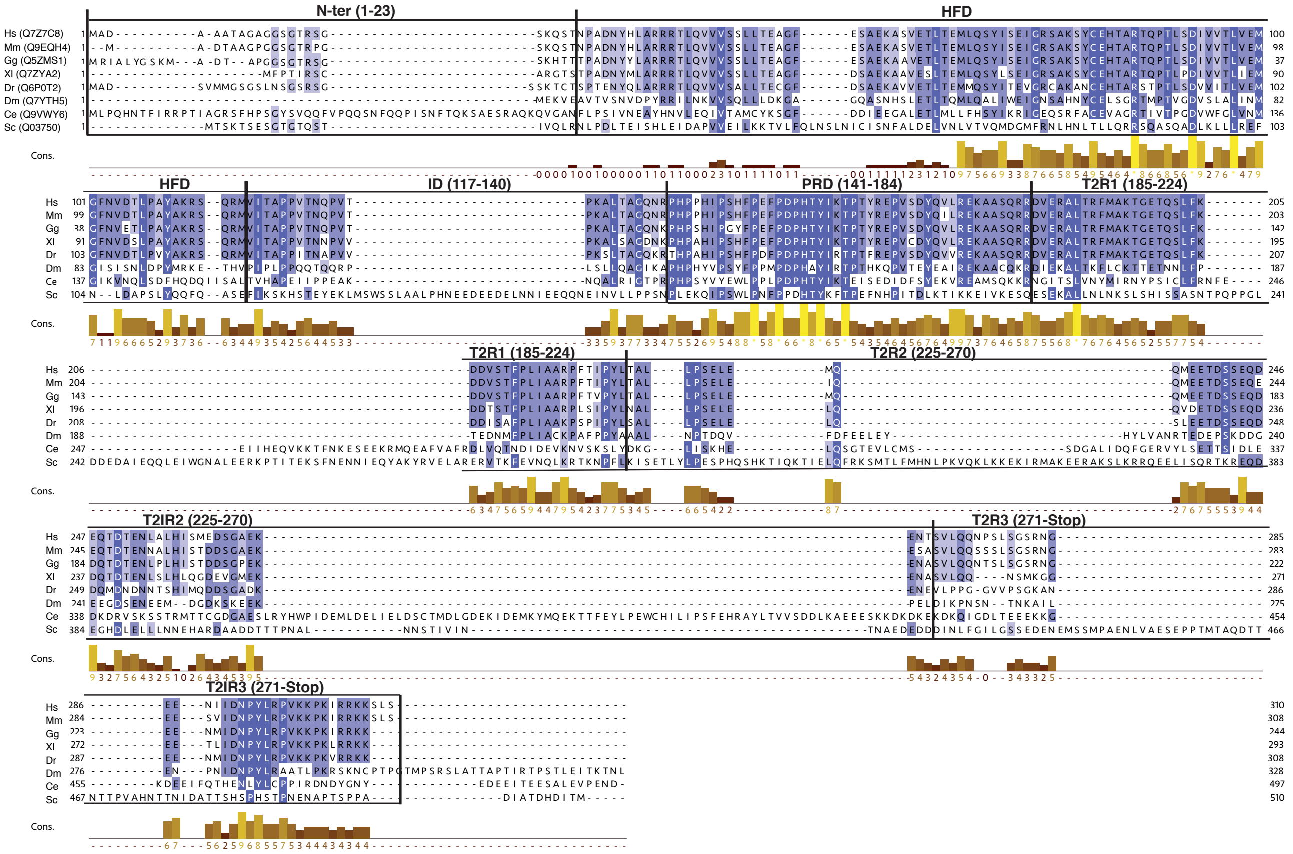

**Figure S1. TAF8 alignments**

Multiple sequence alignments of different TAF8 protein species were made by Jalview (https://www.jalview.org/). Identical amino acids are shaded with the same color. Accession numbers are indicated in brackets at the beginning. Hs: *Homo sapiens*; Mm: *Mouse musculus*; Gg: *Gallus gallus*; Xl: *Xenopus laevis*; Dr: *Danio rerio*; Dm: *Drosophila melanogaster*; Ce: *Caenorhabditis elegans*; Sc: *Saccharomyces cerevisiae*. The different domains of TAF8 used in this study and their borders are indicated. N-ter: N-terminal domain; HFD: histone fold domain; ID: intermediary domain; PRD: proline-rich domain; T2R1-T2R3: TAF2-interacting regions 1-3. Cons.: conservation of total alignment.

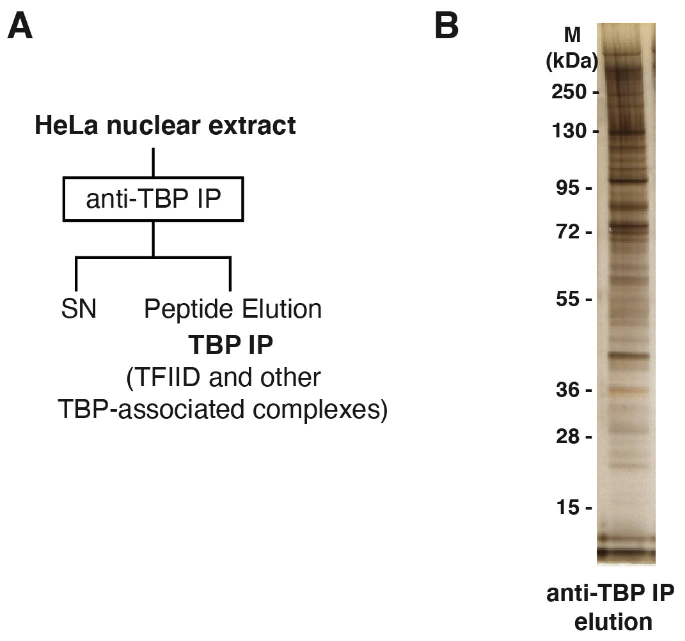


**Figure S2. Anti-TBP IP analysis**

**(A)** Schematic representation of the anti-TBP IP from HeLa cell nuclear extracts using the 2C1 mAb. **(B)** Silver stained gel of the TBP-IP eluted complexes before the CXMS analysis. M: molecular weight markers (indicated in kDa). For the identifications of proteins and CXMS analyses of the IP-ed complexes see Tables S1 and S2.

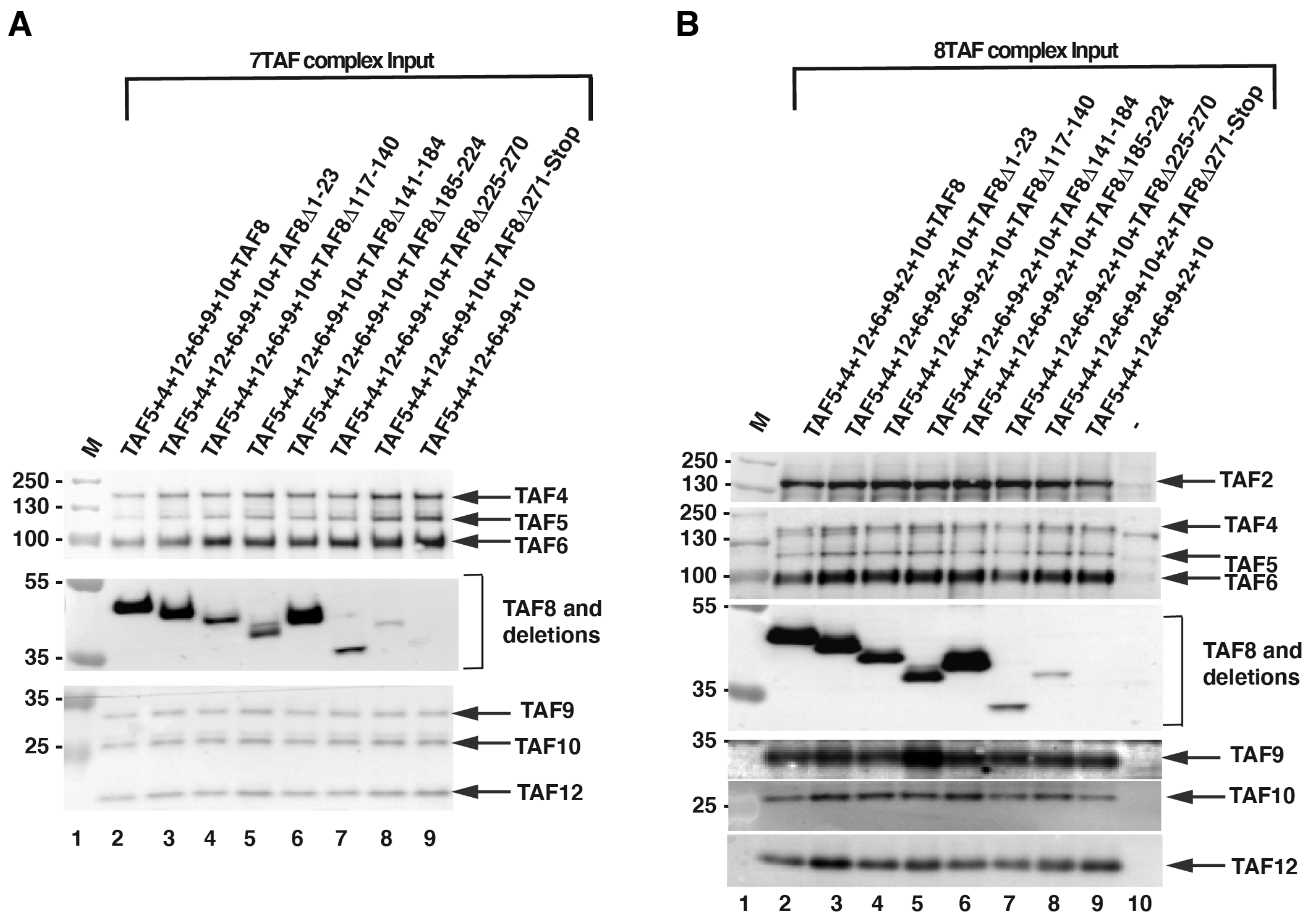


**Figure S3. 7TAF and 8TAF complex coinfections and analysis of the input extracts**

7TAF **(A)** and 8TAF **(B)** complex subunits were co-expressed with either TAF8, with TAF8 deletions or without TAF8, as indicated above each lane, using the baculovirus overexpression system. Input extracts were tested by western blot analyses with the indicted antibodies. Molecular weight markers (M) are indicated in kDa. In (B) - : indicates Sf9 cell extract without any coinfection.

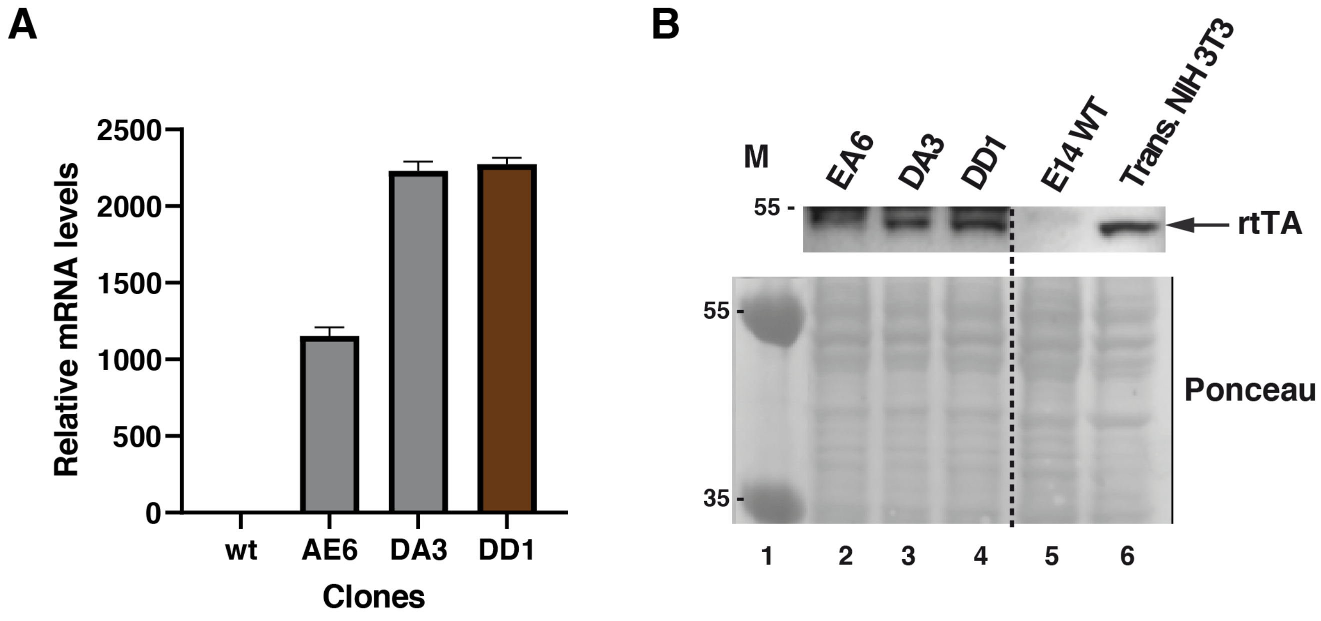


**Figure S4. Establishment of the rtTA expressing E14 mESC line.**

**(A-B)** To generate a Dox-inducible expression mouse embryonic stem cell system, an expression cassette encoding the reverse tetracycline-controlled trans-activator (rtTA; Gossen et al. 1995) was integrated in E14 ESCs (rtTA-ESC). In pre-selected individual E14 ESC clones (AE6, DA3 and DD1) the expression of rtTA was verified by RT-qPCR (A) and western (B) blot assays as described in the Experimental Procedures section. WT: non modified wild type E14 ESC RNA or protein extracts. In (B) the rtTA expression was verified with anti-VP16 antibodies. Transfected (Trans) NIH3T3 cells were used as positive controls. Ponceau staining was used to verify equal loading. Dotted line indicates were the blot was cut. Molecular weight markers (M) are indicated in kDa.
